# *Arabidopsis thaliana* genes contributing to differences in the outcome of infection with generalist and specialist strains of *Turnip mosaic virus* identified by genome-wide association studies

**DOI:** 10.1101/2020.11.25.397661

**Authors:** Anamarija Butković, Rubén González, Mark Paul Selda Rivarez, Santiago F. Elena

**Affiliations:** Institute de Biología Integrativa de Sistemas (I^2^SysBio), CSIC-Universitat de València, Paterna, València, Spain; Department of Biotechnology and Systems Biology, National Institute of Biology, Ljubljana, Slovenia; The Santa Fe Institute, Santa Fe NM 87501, USA

**Keywords:** emerging viruses, GWAS, host-range, *Potyvirus*, specialism-generalism continuum, virus, evolution, virus-host interactions

## Abstract

Pathogens can be classified as generalists or specialists depending on their host breadth. While generalists are able to successfully infect a wide variety of host species, the host range of specialists is limited to a few related species. Even though generalists seem to gain an advantage due to their wide host range, they usually pay a cost in terms of fitness within each host species (*i.e*., the jack-of-all trades, master of none). On the contrary, specialists have high fitness within their own host. A highly relevant yet poorly explored question is whether generalist and specialist viruses differ in the way they interact with their host’s gene expression networks. To identify host genetic factors relevant for the infection of specialist or generalist viruses, we undertook a genome-wide association study (GWAS) approach. Four hundred fifty natural accessions of *Arabidopsis thaliana* were inoculated with turnip mosaic potyvirus strains that were either generalist (TuMV-G) or specialist (TuMV-S). Several disease-related traits have been associated with different sets of host genes for each TuMV strain. While most of the mapped loci were traitor strain-specific, one shared locus was mapped for both strains, a disease resistance TIR-NBS-LRR class protein. Likewise, only one locus was found involved in more than one of the disease-related traits evaluated, a putative cysteine-rich receptor-like protein kinase 20. To validate these results, the corresponding null mutant plants were inoculated with TuMV-G or -S and the outcome of infection was characterized.

**Author summary:** Generalist and specialist viruses are commonly found in nature, where they have potential for epidemics, and are classified depending on their host breath. In this study we used a genome-wide association study to characterize differences in the genetic basis of both infection strategies from a host perspective. Our experimental setup consisted of 450 accessions of *A. thaliana* and two strains of TuMV. We found differences in the number of associated genes and their functions in disease-related traits. Results were validated by characterization of viral infections in null mutant plants deficient for a set of the identified genes.

## Introduction

Viruses are being constantly challenged by multiple factors, including heterogeneity in their host species, or even differences in susceptibility among genotypes within the same host species. Some viruses adapt to a given host (genotype or species) and are able to replicate in it efficiently. These viruses are called specialists. Specialist viruses pose a great threat *e.g*. to monocultured crops since well-adapted viruses usually show enhanced within-host replication rates often associated to stronger symptoms. Our most important crops are monocultures and specialist viruses cause significant loses every year [1-3]. Examples of specialist viruses are Dengue and mumps from mammalian viruses and barley stripe hordeovirus from plants [2,4]. Other viruses infect hosts from widely different genotypes, species, or even higher taxonomical units, and are called generalist viruses [4]. Cucumber mosaic cucumovirus (that infects more than 1000 plant species) and influenza A virus (that infects birds, humans and other species of mammals) are examples of generalist viruses [4].

By specializing in a single host, a virus can limit interspecific competition and better access limited resources [4]. It is proposed that selection favors specialist viruses because there is a trade-off limiting the fitness of a generalist virus in any of the alternative hosts and evolution proceeds faster in narrower niches [5]. The advantage of a virus being a generalist is that they are able to replicate in multiple hosts. There are limitations to generalism, such as by being able to infect multiple hosts a virus does not maximizes fitness in any particular host [4], confirming the *jack-of-all-trades is a master of none* hypothesis [6]. Some of the mechanisms behind this could be related to antagonistic pleiotropy, where beneficial adaptations to one particular host could be disadvantageous in another [7]. Also, to infect multiple hosts, viruses might need to encode for additional genetic information that slows down its replication. Thus, being simultaneously exposed to different host environments viruses cannot adapt to only one [8,9], thus jeopardizing their evolvability. Also, mutations that are fixed and compensate for antagonistic pleiotropy limit access to different evolutionary paths to global maxima in the fitness landscape [10]. All this makes specialists capable of faster evolution and adaptation than generalists in the face of perturbations or new environments [11]. This is important because climate change leads to new environments and expansion of species into new zones, which could lead to novel hosts for certain viruses that are indigenous for that zone. There are also viruses that can spread to new zones via vectors or plant trade and cause significant economical loses in the indigenous plant species because there would be no preparation for the virus emergence.

Both generalists and specialists can be emerging viruses in naïve hosts that have not developed any form of immunity against the new pathogen. New emerging viruses can originate epidemics or pandemics with serious repercussions to human and animal health or agricultural production. Another serious problem of virus emergence is that even without causing epidemics/pandemics and leading to species extinction they can affect the diversity, density and productivity of the host thus disturbing the equilibrium of the ecosystem [12,13]. This situation should be avoided as greater species-diversity means more productivity, resilience to pathogens and faster recovery from perturbations in the ecosystems [14]. The understanding of the early evolutionary and genetic factors in infectious disease emergence, may help to predict new emerging pathogens and protect important agroecological resources [15]. RNA viruses are a major threat as emerging pathogens mainly because of their high replication and mutation rates, large population sizes and short generation times which leads to high evolvability [16-20].

Genome-wide association studies (GWAS) could help to identify host genetic factors that respond to a pathogen infection. This method has gained popularity over the last 20 years due to the increasing number of genome sequences available for a wide range of organisms [21,22]. The basis of GWAS is capturing the single-nucleotide polymorphisms (SNPs) along the genome of an organism and, using statistical methods (such as linear mixed models), to infer the association of an SNP to the trait being analyzed. The common disease-common variant hypothesis posits that common interacting alleles at multiple disease-predisposing loci underlie most common diseases [21]. This hypothesis would justify the use of GWAS to identify the alleles connecting phenotypes with genotypes. This connection permits identification of genetic risk factors for disease, such as susceptibility and resistance to viral infections [23]. One of the most relevant inferences from GWAS is trait heritability, which indicates how much of the observed phenotypic variation is explained by genotypic variation or the genotyped SNPs relative to the contribution of environmental factors [24]. Therefore, heritability can help in understanding the observed phenotype and its underlying genetic complexity.

Identifying host factors responsible for resistance or permissiveness to infection is essential in the study of pathogens as it may be crucial for disease management. To identify some of these host genetic factors, this work studied the infection of generalist and specialist strains of *Turnip mosaic virus* (TuMV; genus *Potyvirus*, Family *Potyviridae*) in 450 natural accessions of *Arabidopsis thaliana* (L.) Heynh. TuMV infects mostly *Brassicaceae* and is widespread worldwide causing huge economical loses by damaging important crops and other plant species [25,26]. The viral strains used in this study were obtained by Navarro et al. [27] by experimentally evolving a naïve ancestral TuMV isolate in *A. thaliana* mutants deficient in different disease signaling pathways or in presence of recessive susceptibility genes. Among all the resulting viral lineages, one evolved in the *enhanced disease susceptibility 8* (*eds8-1*) mutant, hereafter referred as TuMV-G, and another one evolved in the *jasmonate insensitive 1* (*jin1*) mutant, referred as TuMV-S, showed strikingly different host ranges. The *eds8-1* plants lacked the EDS8 protein, causing the reduction of the expression of plant defensin genes and reduced induced systemic resistance (ISR) but enhanced systemic acquired resistance (SAR). The *jin1* plants lacked the JIN1 protein, causing the loss of jasmonic acid (JA) signaling which is a negative regulator of salicylic acid (SA)-dependent signaling. This results in a constitute expression of SAR. The *eds8-1* plants turned out to be the most resistant ones to TuMV infection while the *jin1* plants were the most susceptible ones. This difference in susceptibility gave rise to the two TuMV strains, the generalist TuMV-G and the specialist TuMV-S. TuMV-G was able of infecting with equal fitness all tested plant genotypes, while TuMV-S did so only for its local host genotype. Indeed, Navarro et al. [27] calculated Blüthgen’s *d*’ specialization indexes [28], finding that TuMV-G had *d*’ = 0 (no specialization) while TuMV-S had *d*’ = 1 (complete specialization). In agreement with previous potyvirus-*A. thaliana* studies [29,30], more permissive hosts (here *jin1*) select for more specialized viruses while more restrictive hosts (in this case, *eds8-1*) select for more generalist viruses.

One of the natural hosts of TuMV is *A. thaliana* [31]. This plant is undoubtedly one of the most suitable organisms for GWAS. It has over 1000 natural accessions genotyped and described so far from Eurasia, North America and North Africa [32]. It is self-fertilizing and of small size, making the GWAS analyses much easier since you can maintain a genotype by self-fertilization for an unlimited amount of time and phenotype it repeatedly [23]. We chose a GWAS approach to discover host genes that can be involved in a different response to the generalist and specialist TuMV strains, guiding us to the discovery of potential resistance or pathogenicity factors involved in the infection.

In summary, the response to infection of 450 *A. thaliana* natural accessions from different geographic regions was phenotyped in a controlled setting. The accessions were inoculated with TuMV-G or TuMV-S, two viral strains that differ in their degree of specialization. Infection data (disease progression, infectivity and symptoms severity) was analyzed using GWA, where the identified loci specifically associated with the generalist and specialist TuMV infection. The genetic structure of the phenotyped traits was also studied thanks to the tool Bayesian sparse linear mixed model (BSLMM).

## Materials and methods

### Plant material and growth conditions

Four hundred and fifty *A. thaliana* accessions (File S1) from the 1001 *Arabidopsis* genome collection (https://1001genomes.org; 1001 Genomes Consortium, 2016) were phenotyped. The accessions were randomly selected in an effort to represent heterogenous geographical regions. The 450 accessions were grown in growth chambers under long day regime (16 h day/8 h night) with temperature of 24 °C day/20 °C night, 45% relative humidity and 125 μmol m^−2^s^−1^ of light intensity (1:3 mixture of 450 nm blue and 670 nm purple LEDs).

### Virus inoculum and inoculation procedures

The two strains of TuMV used in this study, TuMV-G and TuMV-S were obtained by evolving, during 12 experimental passages, an ancestral TuMV strain in mutant plants of the *A. thaliana* Col-0 accession, as detailed in [27] and summarized in the Introduction.

The virus-infected plant tissue was frozen in liquid N2 and homogenized and prepared with ten volumes of inoculation buffer (50 mM KH2PO4 pH 7.0, 3% polyethylene glycol 6000, 10% Carborundum) right before the mechanical inoculation. After that, the two TuMV strains were mechanically inoculated into healthy *A. thaliana* plants of the corresponding accession that were between 21 – 25 days old. The inoculation started from the plants that were the largest (8 – 12 leaves) giving the smaller plants extra time to grow so all the accessions got inoculated at a similar size (growth stage ~3.5 in Boyes’ scale [33]). Three leaves were mechanically inoculated with 5 μl of infectious sap prepared in inoculation buffer.

Eight plants per accession for each TuMV strain were inoculated, resulting in a total of 16 plants phenotyped and two mock-inoculated control plants per accession. The accessions were split into two blocks because of chamber space and workforce capacity. The inoculation procedure took about 3 – 4 days. First block was inoculated from 2019/05/06 to 2019/05/08 and the second block was inoculated from 2019/09/11 to 2019/09/14. To reduce spatial correlations due the position of each plant in the growth chamber, pots were translocated to a new random position every day.

### Phenotyping

Three phenotypic traits were measured: (1) Disease progression, calculated as the area under the disease progression stairs (*AUDPS*) [34]. The number of infected plants was quantified daily and the figures used to calculate the progression of the disease through time. (2) Infectivity: number of infected plants out of the total number of inoculated plants after 21 days post-inoculation (dpi). (3) Symptoms severity: on a scale from 0 – 5 (Fig. 1) measured at intermediate (14 dpi) and late (21 dpi) infection times to explore time-dependent differences in gene expression. As previously described [30,35], the presence and strength of symptoms of TuMV in *A. thaliana* natural accessions is directly correlated with viral accumulation.

**FIG 1.**
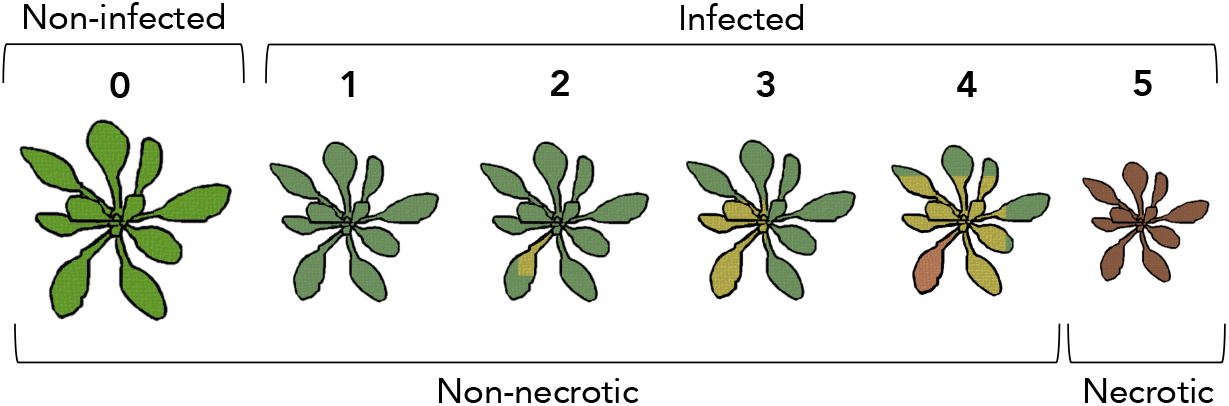
Symptoms scale that was used to evaluate the severity of symptoms in the plants during the 21 days period post inoculation. 0: no symptoms or healthy plant, 1: mild symptoms without chlorosis, 2: chlorosis is visible, 3: advanced chlorosis, 4: strong chlorotic symptoms and beginning of necrosis, 5: clear necrosis and death of the plant.

In the characterization of plant knock-out (KO) null mutant response to infection, *AUDPS* and symptoms intensity progression steps curve (*AUSIPS*) [36] were measured. *AUSIPS* is calculated using daily symptoms intensity values and, similar to *AUDPS* that summarizes disease progression, it summarizes the progression of the symptomatology through time.

### Genome-wide association mapping

Association analyses were done with a Python program based on LIMIX [37] written by Prof. Magnus Nordborg’s group. The kinship matrix and the genotype data come from the 1001 Genome project in *A. thaliana* [32], consisting of the SNPs for the 1135 genome accessions plus imputed SNPs of a set of accessions that were genotyped with a 250k SNP chip. In LIMIX, SNPs and covariates are treated as fixed effects while the population structure and noise are treated as random effects in the linear mixed model.

The normality of the data was checked with SPSS version 25 (IBM Corp., Armonk NY, USA). Untransformed phenotypic data were used in the GWAS. Since phenotyping was done in two blocks (150 accessions + 300 accessions), block effects were accounted for in the GWAS analysis.

Out of ~10 million SNPs [38], 1,815,154 SNPs had a minor allele frequency higher than 0.05 for all phenotypes. The multiple testing problem is rooted in the assumption that each test performed is independent of others, which is often not true due to linkage disequilibrium between genetic markers. This can lead to false negatives and important genes related to our phenotype might not be discovered [21,23]. Therefore, the false discovery rate (FDR) was used to deal with false positives and false negatives. The FDR was calculated using the fdrtool package version 1.2.15 in R version 3.6.1 in RStudio version 1.2.1335. The exact FDR values used were as follows: for TuMV-G *AUDPS* 21 dpi FDR = 2.73×10^−10^, infectivity 14 dpi FDR = 6.32×10^−13^ and 21 dpi FDR = 1.78×10^−8^, and symptoms 14 dpi FDR = 9.59×10^−10^. While for TuMV-S it was calculated only for symptoms trait at 14 dpi FDR = 1.49×10^−12^ and 21 dpi FDR = 1.15×10^−9^. If FDR could not be calculated, −log*P* ≥ 5 was used as a conservative threshold. This was the case for *AUDPS* 14 dpi and symptoms 21 dpi in the case of TuMV-G and *AUDPS* 14 and 21 dpi for TuMV-S. Manhattan and quantile-quantile (QQ) plots were drawn using rMVP package [39] in R version 3.6.1 in RStudio version 1.2.1335.

### Bayesian sparse linear mixed model (BSLMM)

BSLMM implemented in GEMMA was used to infer the genetic architecture of the measured phenotypic traits [40,41]. With this method it was determined which loci show the strongest correlation with the trait in question and the genetic basis of the trait: whether it is highly polygenic or rather determined by a few loci (*i.e*., oligogenic). Raw values were used for symptoms severity (discrete variable) while *AUDPS* and infectivity (continuous variable) were normalized by block using a univariate general linear model in SPSS. In all cases, MCMCs were run with the default settings (burn-in at 100,000, sampling steps at 1,000,000 and recording every ten steps) and minor allele frequency cut-off set at 5%.

The proportion of variance in phenotypes explained (*PVE*) by available genotypes, or total variance explained by additive genetic variants, was used as an estimator of the heritability of a given phenotypic trait. *PVE* is a flexible Bayesian equivalent of the narrow sense heritability (*h*^2^) estimated by more classical linear mixed models (LMM). In addition, BSLMM estimated the proportion of genetic variance explained by sparse effects (*PGE*), its maximum value being *PGE* ≈ 1 for highly heritable traits.

### Validation of GWAS associations

Ten genes identified with the GWAS were selected for further study. Mutant plants with a disruptive insertion in those genes were ordered from the NASC stock center (https://arabidopsis.info/BrowsePage). The mutants were chosen on the following two criteria: (*i*) must be in the Col-0 background, (*ii*) must consist of a T-DNA insert that results in KO of the gene. The mutant plants were seeded on the 2020/06/03 and inoculated as previously described with the two TuMV strains on the 2020/06/23. The *AUDPS* and the *AUSIPS* were calculated using the number of infected plants and their symptomatology measured during 21 dpi for each individual plant. For the values of *AUDPS* and *AUSIPS* a thousand pseudo-replicated matrices, of equal dimensions to the original one, were obtained per experimental condition. The matrix rows (each individual plants infection history) were replaced and thus the temporal correlations across time points were preserved. This algorithm, implemented in R version 3.6.1, generated kernel distributions for *AUDPS* and *AUSIPS*. The highest density intervals (HDI) were calculated using the bayestestR package in R version 3.6.1 in RStudio version 1.2.1335, applying its default value of 89% HDI [42].

## Results

### Characterization of infection traits in natural accessions

The 450 *A. thaliana* accessions were phenotyped for the infection with generalist (TuMV-G) and specialist (TuMV-S) strains of TuMV (S1 Appendix). Several infection-related traits were characterized by visual inspection, one of them being the progression of the disease through time (*AUDPS*). *AUDPS* is an important trait used to characterize the infection since in natural accessions it correlates with virus accumulation and severity of symptoms, as it has been previously described [30,35]. Two-sample Kolmogorov-Smirnov tests were performed to compare the distributions of TuMV-G and TuMV-S *AUDPS* values in the 450 natural accessions (Fig. 2). When compared to the specialist, the generalist virus performed better in a larger set of accessions. This happened at intermediate and late infection times, as TuMV-G showed a significantly different distribution at 14 dpi (*D* = 0.1644, *P* < 0.0001) and 21 dpi (*D* = 0.1622, *P* < 0.0001). Analyses were performed at different infection times as it is known that the set of genes differentially expressed depends on the stage of infection [35]. These temporal differences were corroborated using a two-sample Kolmogorov-Smirnov tests for the distribution of the *AUDPS* values for each strain at both infection times. The test showed that the distributions are significantly different at 14 dpi and at 21 dpi, for both TuMV-G (*D* = 0.9378, *P* < 0.0001) and TuMV-S (*D* = 0.8978, *P* < 0.0001). Therefore, the genome-wide associations of the infection traits were studied at intermediate (14 dpi) and late (21 dpi) infection times.

**FIG 2.**
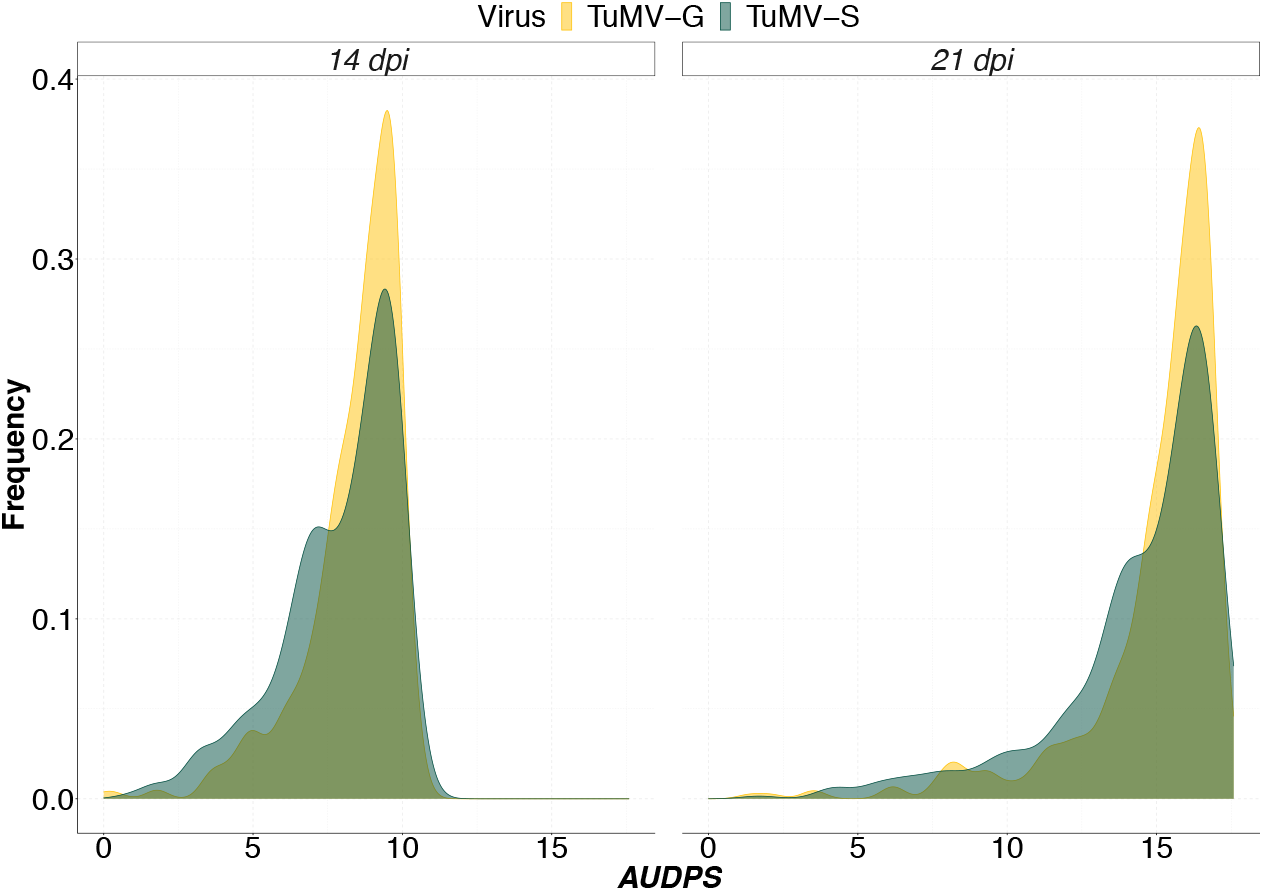
Density plots for TuMV-G (yellow) and TuMV-S (green) *AUDPS* values in the 450 i natural accessions at 14 (left) and 21 dpi (right).

### Genetic architecture of disease-related traits

The 450 accessions accounted for 431,323 SNPs that were tested in both viral strains at both 14 and 21 dpi (S2 Appendix) using the BSLMM analysis. This analysis evaluates how much of the phenotype is explained by the genotyped SNPs in our traits, defined as phenotypic variance (*PVE*).

*AUDPS*, infectivity and symptoms severity had low *PVE* values (S2 Appendix). The lowest *PVE* values were obtained for the traits *AUDPS* for TuMV-G at 14 dpi, and for infectivity and symptoms severity for both strains at both 14 and 21 dpi. Overall, the values were similar for both TuMV strains and between the two time points for *PVE* values (except for *AUDPS* trait at 14 dpi where TuMV-S had higher values than TuMV-G) and the number of major effect variants that explain phenotypic variance (or *PGE*) (S2 Appendix).

In order to detect the large effect SNPs that contribute the most to our phenotype, posterior inclusion probability or the strength of association of a SNP with a phenotype above or equal to 0.25 were used. The percentage of *PVE* explained by large effect variants (*PGE*) indicates that major effect loci explain between 50 - 90% of phenotypic variance in *AUDPS* and infectivity traits, in both time points for both viruses. The number of variants with large effect size, the SNPs that explain most of the phenotype, among the 431,323 SNPs was low for infectivity and symptoms at 14 dpi for both viruses and *AUDPS* and infectivity at 21 dpi for both viruses (S2 Appendix). The frequency of a SNP estimated to have a detectable large effect in *AUDPS* for TuMV-S 21 dpi was mapped within the gene encoding for *AT2G04440*, a MutT/Nudix family protein. *AT2G04440* was previously characterized as an important player in the plant immune response [43]. While for infectivity trait for TuMV-G at 21 dpi the most contributing SNP was mapped within locus *AT3G19350*, that corresponds to the gene *MATERNALLY EXPRESSED PAB C-TERMINAL* (*MPC*). And also, position 6,685,977 of an intergenic region on chromosome 3 was mapped as significant for TuMV-G infectivity at 21 dpi. *MPC* is an important translation initiation factor that binds to the VPg and the RNA-dependent RNA polymerase (RdRP) NIb of TuMV, affecting the viral RNA accumulation [44]. Chromosome 3 intergenic position 6,685,977 is between loci *AT3G19290*, which corresponds to the gene *ABA-RESPONSIVE ELEMENT BINDING PROTEIN 4* (*ABF4*), and *AT3G19280*, which corresponds to the gene *FUCOSYLTRANSFERASE 11* (*FUT11*). This noncoding region between these two genes could be a promoter region involved in regulation of the expression of both *ABF4* and *FUT11. ABF4* controls the ABA-dependent stress response. It was previously shown that wheat yellow mosaic potyvirus disturbs the ABA signaling pathway through the interaction between the viral RdRp and the wheat’s light-induced protein TaLIP thus facilitating virus infection [45]. There is no clear description of *FUT11* in plant virus infection, but it is involved in protein N-linked glycosylation where a carbohydrate is added to a protein.

In summary, for the infection caused by the pathogens studied, the genetic architecture of *AUDPS* and infectivity phenotypes is relatively simple. It involves few SNPs for the two viral strains analyzed while also having a detectable large effect SNP that is being responsible for the majority of the observed phenotype. Symptoms severity is genetically more complex and involves many more small effect SNPs. In both viruses all the phenotypes have a similar genetic architecture between the two temporal stages (14 and 21 dpi).

### GWAS identifies genetic loci associated with disease-related phenotypes induced by specialist and generalist viral strains

The SNPs association significance of the infection traits (*AUDPS*, infectivity and symptoms severity) were visualized using Manhattan plots in (Fig. 3). The QQ-plots for infection traits showed no detectable population structure (S1 Fig). Using the FDR or the −log*P* ≥ 5 threshold (Material and Methods, S3 Appendix) of the traits, a total of 55 significant SNPs were identified for TuMV-G and 94 for TuMV-S infection (S3 Appendix). Some of these SNPs were positioned within seven genes for TuMV-G and 13 for TuMV-S infection traits (Table 1 and S4 Appendix).

**Table 1.** Significant genes detected using GWAS for the three disease-related traits during the course of infection with each viral strain.

| **TuMV-G** |  |  |  |  |  |
| --- | --- | --- | --- | --- | --- |
| **Trait** | **dpi** | **Gene** |  | **Description** | **–log *P*** |
| *AUDPS* | 14 | *AT1G57570* | *JAL14* | Jacalin-related lectin 14 | 5.01 |
| Symptoms | 14 and 21 | *AT2G14080* |  | Disease resistance protein (TIR-NBS-LRR class) family | 9.02 |
| Symptoms | 21 | *AT3G21660* | *PUX6* | Plant UBX domain-containing protein 6 | 5.54 |
| Symptoms | 21 | *AT3G46590* | *TRP2* | Telomere repeat-binding protein 2 | 5.09 |
| Symptoms | 21 | *AT4G04540* | *CRK39* | Putative cysteine-rich receptor-like protein kinase 39 | 5.65 |
| Symptoms | 21 | *AT4G32660* | *AFC3* | Serine/threonine-protein kinase AFC3 | 5.22 |
| Infectivity | 21 | *AT5G66750* | *DDM1* | ATP-dependent DNA helicase DDM1 | 8.00 |
| **TuMV-S** |  |  |  |  |  |
| **Trait** | **dpi** | **Gene** |  | **Description** | **–log *P*** |
| Symptoms | 14 and 21 | *AT2G14080* |  | Disease resistance protein (TIR-NBS-LRR class) family | 11.83 |
| Infectivity | 14 and 21 | *AT4G23280* | *CRK20* | Putative cysteine-rich receptor-like protein kinase 20 | 6.52 |
| *AUDPS* | 21 | *AT4G23280* | *CRK20* | Putative cysteine-rich receptor-like protein kinase 20 | 5.73 |
| *AUDPS* | 14 and 21 | *AT2G04450* | *NUDT6* | Nudix hydrolase 6 | 6.45 |
| *AUDPS* | 14 and 21 | *AT3G21980* | *CRRSP27* | MutT/nudix family protein | 6.11 |
| Infectivity | 14 and 21 | *AT4G23280* | *CRK20* | Putative cysteine-rich repeat secretory protein 27 | 5.91 |
| Infectivity | 14 | *AT4G02580* |  | NADH dehydrogenase [ubiquinone] flavoprotein 2 | 5.64 |
| Infectivity | 21 | *AT4G13345* | *MEE55* | Serinc-domain containing serine and sphingolipid biosynthesis protein | 5.32 |
| Infectivity | 21 | *AT4G10130* | *T9A4.1* | DNAJ heat shock N-terminal domain-containing protein | 5.22 |
| *AUDPS* | 21 | *AT3G07470* |  | Transmembrane protein | 5.20 |
| *AUDPS* | 14 | *AT3G12850* |  | COP9 signalosome complex-related / CSN complex-like protein | 5.18 |
| *AUDPS* | 14/21 | *AT5G08650* |  | Translation factor GUF1 homolog | 5.14 |
| *AUDPS* | 14 | *AT2G04430* | *NUDT5* | Nudix hydrolase 5 | 5.02 |

**FIG 3.**
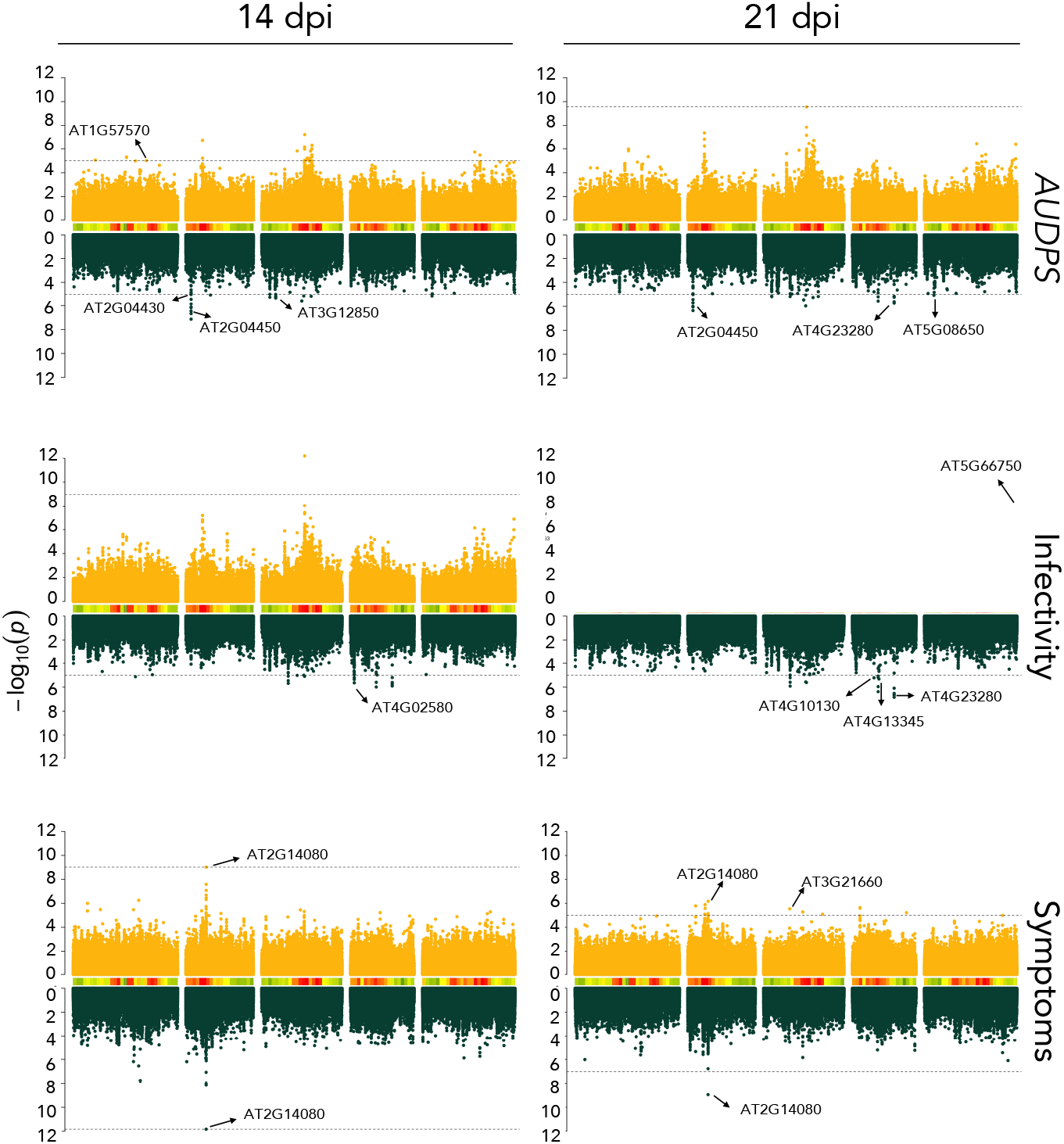
Manhattan plots of the analyzed disease-related traits. Data for TuMV-G are indicated in yellow and for TuMV-S in green. Peaks marked on the plots correspond to the most significant SNP values of the genes selected for the mutant analysis. SNP density shows how many SNPs are genotyped for a particular chromosomal region. The dashed lines indicate the significance threshold (FDR or −log*P* = 5 if it was not possible to calculate the FDR).

Most of the genes were unique for (*i*) the TuMV-G or the TuMV-S strain and (*ii*) the infection trait as seen in Fig. 4. There was one locus shared between the two viral strains, both at 14 and 21 dpi, for symptoms severity: the aforementioned *AT2G14080*. The NBS-LRR genes are the most numerous class of the *R* (resistance) genes in *A. thaliana*. Their effector recognition LRR domains recognize specific pathogens and can lead to a hypersensitive immune response (HR) or to an extreme resistance against the virus infection. An HR restricts the pathogen at the primary infection site causing cell death followed by SAR that increases SA accumulation and expression of pathogenesis-related genes [46,47].

**FIG 4.**
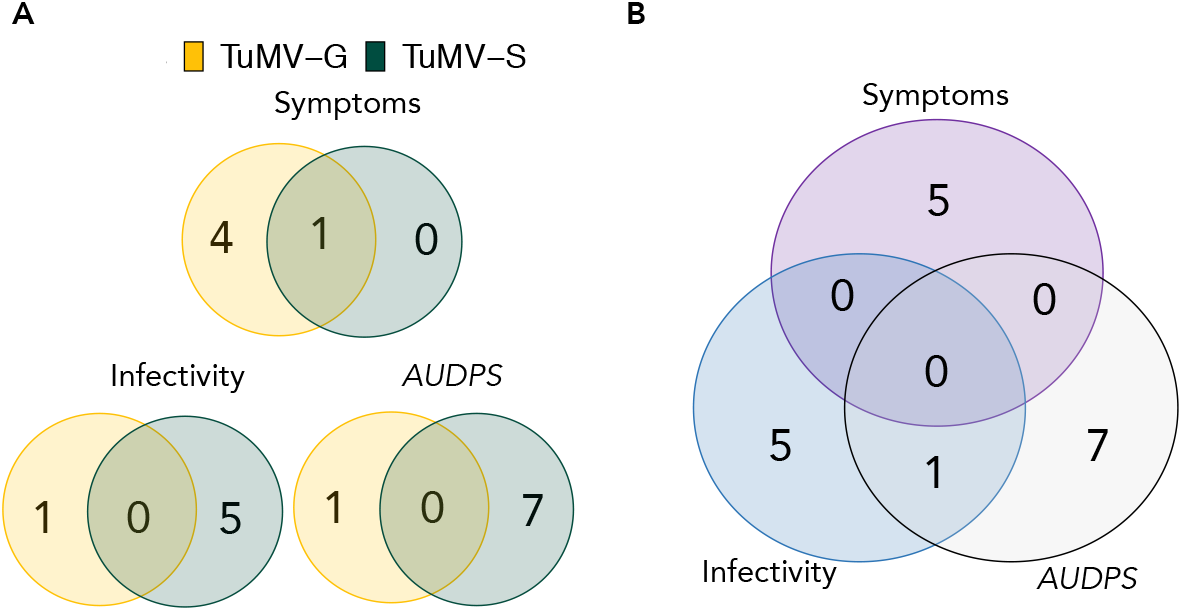
Venn diagram showing the number of unique and shared genes. (A) Genes mapped for each viral strain and disease-related traits. (B) Genes mapped for all disease-related traits pooling together both viral isolates.

Comparing the results at 14 and 21 dpi for TuMV-G, genes at 14 dpi seem related to a more general disease response while the genes at 21 dpi are more specific and involved in ubiquitin-related processes. Such difference is not seen between the 14 and 21 dpi for TuMV-S. This suggest that the generalist strain does show a difference in host gene involvement depending on the stage of the infection while the specialist strain does not.

### Experimental validation of identified genes

Ten of the identified genes were selected for a validation study in which the corresponding KO mutants were inoculated with both viral strains and the disease progression characterized (Fig. 5 and S5 Appendix). Out of the 10 genes, one was shared between the two viral strains, two were unique for TuMV-G and seven were unique for TuMV-S. More genes were validated for TuMV-S because the GWAS mapped more significant SNPs upon infection with this strain. The selected KO mutants were: *at1g57570, nudx5, nudx6, at2g14080, at3g12850, at4g10130, mee55, CPLEPA, ddm1*, and *at4g02580*. To evaluate the difference in the infection dynamics between the mutants and the wild-type (WT) plants, the *AUDPS* and *AUSIPS* values were calculated using the data collected along the 21 dpi. A comparation between the WT and KO mutant values for each viral strain was done (Fig. 5, S6 Appendix) based on the inferred 89% HDIs. Differences in most of the KO were found when comparing the *AUDPS* values of the two viral strains with the WT (S6 Appendix).

**FIG 5.**
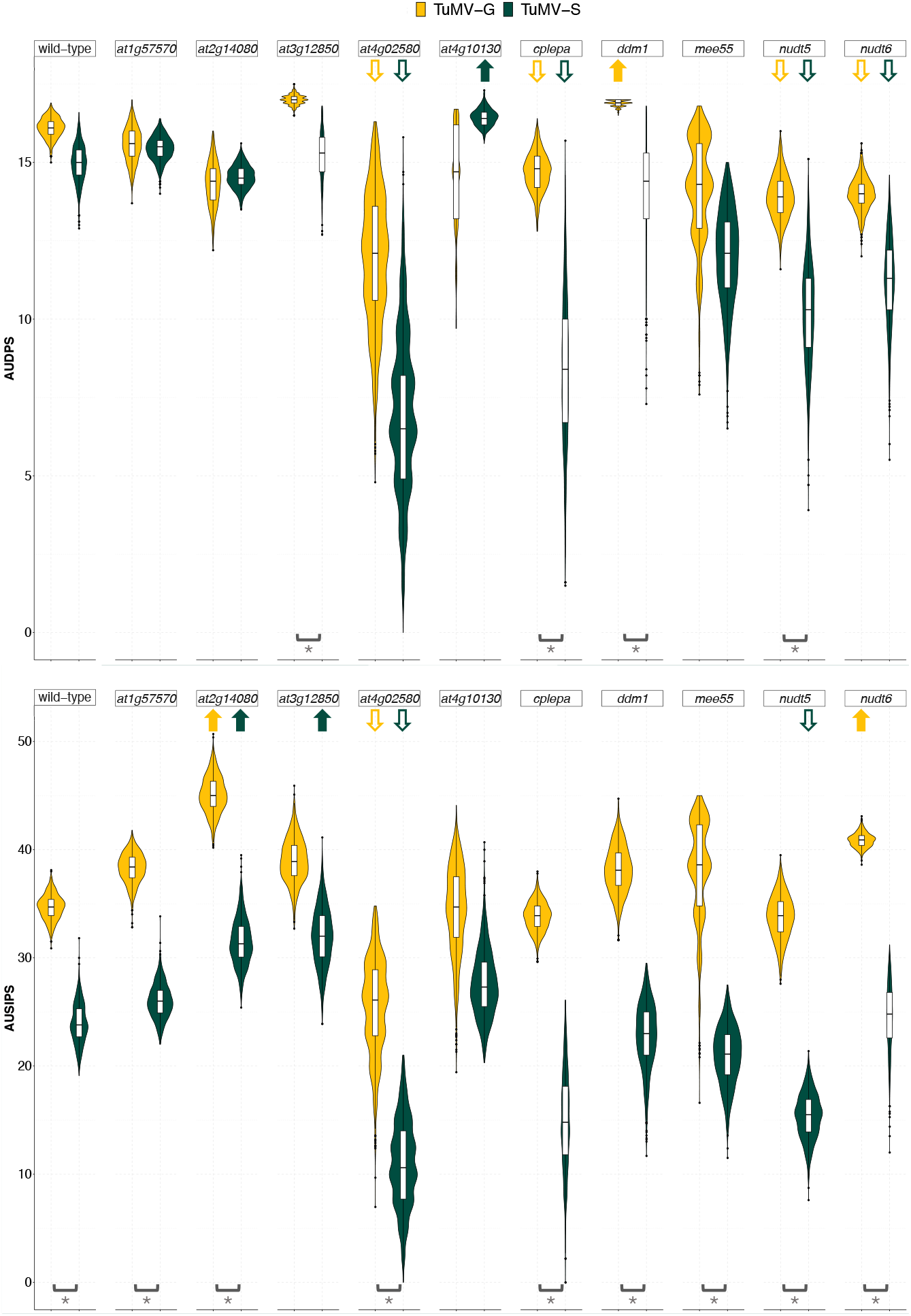
89% HDI calculated for *AUDPS* and *AUSIPS* for each viral strain on each KO mutant plant genotype. Not overlapping 89% HDI between a given mutant and the wild-type plants is indicated by an arrow. Arrows pointing up indicate a significant positive difference in medians, while arrows pointing down indicate the opposite trend. Brackets and asterisks indicate significant differences between TuMV-G and TuMV-S disease progression or severity in the KO mutant plant genotype being considered.

Evaluating the mutant *AUDPS* intervals, lower HDIs compared to the WT imply that these mutants have slower disease progression because the KO gene is positively involved in the viral cycle and the virus uses it to aid its replication or translation. For the TuMV-S and TuMV-G infection, there are four mutants that have lower HDI compared to the WT: *at4g02580, cplepa, nudt5*, and *nudt6. AT4G02580* is a susceptibility factor and could aid viral pathogenesis [48]. CpLEPA is a highly conserved chloroplastic translation factor that could assist viral transcription in the cytoplasm by enhancing the translation of chrolopastic proteins involved in photosynthesis to compensate for the negative side-effects of infection in chloroplasts activity [49-51]. The *nudt5* and *nudt6* are deficient in proteins that form part of the Nudix hydrolase family, which act as positive regulators in plant immunity [43,52]. Seeing how here they have a role as positive regulators in plant immunity, and the opposite was observed in our validation study, there is a possibility that these two Nudix hydrolases could have other undescribed functions.

Observing KO mutants with *AUDPS* intervals higher than the WT suggests that the corresponding genes are involved in the defense response of the plant to infection, since removing them enhanced disease progression beyond that observed for the WT plants. In the TuMV-G infection, *ddm1* had HDI higher than the WT. Corrêa et al. [35] showed that *ddm1* plants were more resistant to two different strains of TuMV. This might be because induction of SA-mediated defense in the *ddm1* mutants may be an explanation of their resistance to the virus. The opposite has been noticed for geminiviruses where the *ddm1* mutant showed hypersusceptibility to infection. The reason for this was the methylation of viral genomes which is a defense mechanism of the plant and when the methylation is reduced the plants are more susceptible [53]. The different adaptation history of TuMV-G compared to the strains studied by Corrêa et al. [35] might explain this discrepancy. For TuMV-G the lack of DDM1 might help the virus replicate better since defense genes are not properly methylated and henceforth their expression deregulated. In plants infected with TuMV-S only *at4g10130* had higher 89% HDI compared to the WT. *AT4G10130* is involved in peptidyl-diphthamide biosynthetic processes and tRNA wobble uridine modification. Both of these processes are involved in translation modifications and this protein might have a role in an anti-pathogenic response.

For the *AUSIPS*, if mutant values have higher intervals than the WT it will mean that the virus is able to cause stronger symptoms in the absence of these host genes. This is the case for the *at2g14080* and *nudt6* plants infected with TuMV-G. In the case of TuMV-S this happened in *at3g12850* and, as for TuMV-G, in *at2g14080* plants. *AT2G14080* encodes for a disease resistance protein as described before. These proteins monitor the status of plant proteins targeted by pathogens and activate a series of defense responses [54]. By removing this gene, viruses managed to cause stronger symptoms. NUDX6 acts as a positive regulator of the SA signaling pathway by modulating NADH levels [52]. This leads to a higher pathogen response and KO mutant *nudt6* allows the virus to replicate better giving it higher credible intervals compared to the WT. An interesting observation was made for this mutant in the *AUDPS* measurement, where it showed an opposite effect in comparation with *AUSIPS. AT3G12850* is involved in regulation of JA levels. Viruses infecting *at3g12850* plants replicate better. *at3g12850* is a COP9 signalosome complex-related/CSN complex-like protein. The tomato yellow leaf curl Sardinia geminivirus (TYLCSV) C2 protein interacts with CSN5 resulting in a reduction of JA levels. As previously shown, treating *A. thaliana* plants with exogenous JA disrupts TYLCSV infection [55]. It is known that plant viruses and herbivores have strategies to manipulate plant JA levels as this hormone confers defenses to the plant against biotic and abiotic stresses [56]. This means that in our pathosystem the JA is negatively affecting the viral replication.

Therefore, significant differences in *AUDPS* intervals between WT and mutant plants confirm the role in infection of the genes that were knocked-out. When in this comparation the mutant had lower *AUDPS* values (*e.g., at4g02580, cplepa, nudt5*, and *nudt6* for both viral strains) it confirms the function of the gene in the viral replication. While mutants with values higher than those of the WT (*e.g., ddm1* mutants for TuMV-G and *at4g10130* for TuMV-S) confirm the role of the gene in the host defense. Comparations of *AUSIPS* values between WT and mutant plants also confirms the role of most of the studied genes in symptoms severity. The *at4g02580* plants had lower *AUSIPS* interval in the TuMV-G infection. Therefore, plants defective in a NADH-ubiquinone oxidoreductase susceptibility factor had a milder symptomatology than WT ones. For the infection with TuMV-S, *at4g02580* and *nudt5*, also had lower *AUSIPS* intervals. The difference in symptoms severity progressions was also significant between the two viral strains in mutants WT, *at1g57570, at2g14080, at4g02580, cplepa, ddm1, mee55, nudt5*, and *nudt6*. This difference indicates that the two viral strains cause different symptomatology in the WT and the majority of mutants.

## Discussion

Pathogens will have different virulence and induce different responses in the hosts they are infecting depending on whether they are generalists or specialists. For example, in the whole-genome transcriptomic study by Hillung et al. [57] they compared the transcriptomic responses of six *A. thaliana* accessions infected with generalist or specialist tobacco etch potyvirus (TEV) strains. They showed that the generalist virus manipulates a similar set of host genes across the host range while the specialist virus shows a more heterogeneous response. In the GWAS study here presented, similar conclusions have been reached by comparing genes involved in the infection of *A. thaliana* with generalist or specialist TuMV strains. In the case of the generalist strain TuMV-G, fewer candidate genes were identified than in the case of the specialist strain TuMV-S. This difference emerged as a consequence of the different evolutionary strategies of the two viruses. Selection has driven the generalist virus to manipulate a similar set of host genes across the host range for successful infection. Therefore, the GWAS detected a limited number of candidate genes. In contrast, host-specific selective pressures modulated the evolution of the specialist virus, henceforth, more genes associated with TuMV-S have been found by the GWA analysis.

Mapped genes in the GWAS analysis belonged to categories such as F-box proteins, kinase, hydrolase, LRR family proteins, disease resistance proteins, transcription factors, lectins, helicases, ubiquitin proteases, proteins involved in iron metabolism, pentatricopeptide repeat-containing, GTPases, and berberines. The locus *AT2G14080* was identified in common for both viral strains in the analysis of symptoms severity. There were also some strain-specific hits that were previously characterized as involved in plant defense or in some important part of the viral cycle. Indeed, genes that differ between the two viruses could be targets of differential selection for specialist or generalist viruses to evolve. For example, ubiquitin protease and TELOMERE REPEAT-BINDING PROTEIN 2 were specific responses to TuMV-G infection. While the Nudix hydrolase, NADH dehydrogenase and DNAJ heat shock proteins were specific of plants infected with TuMV-S strain.

In the analysis of the ten selected KO null mutants significant differences can be detected with *AUDPS* and *AUSIPS*, indicating that disease progression was not proportional to symptoms development in the mutants. The reason for this effect could be that the viral load in the a given Col-0 mutant, as opposed to the natural accessions, is not proportional to symptoms severity. Symptom appearance and progression depend on the viral load but symptom severity might depend on the lack of an essential gene the virus might hijack to evade the defense response, not being directly related to viral load. The virus not being able to evade the defense response might activate stronger SAR which leads to stronger symptoms appearance because of the stronger hypersensitive immune response which restricts the pathogen at the primary infection site causing cell death. There was one gene that came up in the GWAS of both strains, *AT2G14080* that had a significant effect in the mutant involved with the two strains and it appears to be involved in plant defense. Two of the ten genes selected for the mutant analysis came from TuMV-G analysis and seven came from the TuMV-S analysis. In total out of the ten selected genes, eight had a significant effect on the virus disease progression and/or symptoms. *MEE55* (a serine and sphingolipid biosynthesis protein, MATERNAL EFFECT EMBRYO ARREST 55) and *AT1G57570* (a member of the mannose-binding lectin superfamily protein) apparently had no significant effect on either viral strain under our conditions. There were five genes that had an effect in both viral strains: *AT2G14080, AT4G02580, cpLEPA, NUDX5*, and *NUDX6*. Nudix hydrolases 5 and 6 are involved in plant defense [52] but in both strains they seem to have important roles for disease progression by enhancing viral replication or gene expression since the viruses replicate worse when these two genes where KO. This role for the two hydrolases in viral infection was not described before.

A GWAS of TuMV infection in *A. thaliana* in a natural setting was performed by Rubio et al. [58]. None of the genes found by these authors was pinpointed in our study but this could simply reflect three major experimental differences: (*i*) Rubio et al. grew their plants in a natural setting where they were exposed to a changing environment. The highly complex natural setting can lead to much more heterogeneous gene regulations, as opposed to a controlled environment that minimizes external abiotic and biotic stressors. It was shown before that differences in temperature, light and water availability influence the response of the plant to a virus [59,60]. Multiple stresses affecting the plant at the same time can be problematic when trying to identify genes responsible for the specific response of plants to virus infection. (*ii*) The evolutionary histories of the TuMV strains used in both studies were largely different. While Rubio et al. used the UK1 isolate, we used generalist and specialist strains derived from the YC5 isolate originally obtained from calla lily plants [61]. (*iii*) In our study, the 450 accession were randomly chosen to represent the world-wide genetic diversity of the species, while French accessions were largely overrepresented in Rubio et al. study. Rubio et al., 2019 identified six new genes above a threshold of −log*P* ≥ 4 in their GWAS analysis: *RESTRICTION TO TOBACCO ETCH VIRUS MOVEMENT 3*, a DEAD box RNA helicase 1 candidate gene, *EUKARYOTIC TRANSLATION INITIATION FACTOR 3B*, a protein with a pleckstrin homology domain, a protein containing a TIM barrel domain, and a key enzyme involved in the glutamate pathway. Our study identified 13 genes specifically mapped for viral infection response (Table 1), of which eight were experimentally confirmed as having roles in the plant response to TuMV-S and TuMV-G (Fig. 5). Despite there not being common mapped genes between both studies, there are similarities at the functional level: ATP-dependent DNA helicase, DnaJ domain superfamily protein and ubiquitin associated proteins.

Looking at the analysis of the underlying genetic architecture of each phenotyped trait, it was evident that some disease-related phenotypes were explained by few SNPs (infectivity and symptoms severity at 14 dpi for both viruses and *AUDPS* and infectivity at 21 dpi for both viruses as well), while some traits were highly polygenic and explained by a large number of SNPs (*AUDPS* for TuMV-S at 14 dpi). SNPs that passed the posterior inclusion probability threshold were mapped within locus *AT2G04440* (MutT/Nudix family protein) for *AUDPS* of TuMV-S at 21 dpi and position 6,685,977 in an intergenic region on chromosome 3 for infectivity of TuMV-G at 21 dpi. All had possible roles in the viral infection.

Since the genome of *A. thaliana* is highly polygenic and is governed by small effect loci (as seen in the BSLMM analysis) our study might have missed some of the genes described in the literature as being involved in the potyvirus infection. Other explanation would be that these genes were not important in the context of our virus strains that were preadapted in specific mutants of *A. thaliana*.

Altogether, this work (*i*) describes differences between a generalist and a specialist pathogen, (*ii*) identifies and characterizes genes involved in a generalist and a specialist virus infection and (*iii*) illustrates the variability of the genetic elements involved in a viral infection depending on the evolutionary history of the viral strain.

## Acknowledgements

We thank Paula Agudo and Francisca de la Iglesia for excellent technical support, Prof. Magnus Nordborg’s group for providing the seeds of *A. thaliana* accessions and the Python code for the GWAS. Work was funded by Spain’s Ministerio de Ciencia e Innovación-FEDER grant PID2019-103998GB-I00 and Generalitat Valenciana grants GRISOLIAP/2018/005 and PROMETEU2019/012 to SFE. RG was supported Ministerio de Ciencia e Innovación-FEDER contract BES-2016-077078. MPSR was supported by a Young Investigators fellowship from the Universitat de València International 0.7 Cooperation Program.

## Supporting information

**S1 Fig. The QQ-plots for the phenotyped traits.** (PDF)

**S1 Appendix. The selected accessions.** (EXCEL)

**S2 Appendix. Results of the BSLMM analysis on the four disease-progression selected traits and their 89% HDI**. (EXCEL)

**S3 Appendix. Significant SNPs in the GWAS analysis of TuMV-G and TuMV-S**. (EXCEL)

**S4 Appendix. Significant genes in the GWAS analysis of TuMV-G and TuMV-S with thresholds used and comparation between strains with duplicate hits removed**. (EXCEL)

**S5 Appendix. Selected KO mutants with a description of the corresponding genes functions and a link to reference**. (EXCEL)

**S6 Appendix. 89% HDIs calculated for *AUDPS* and *AUSIPS* for both viral strains on each KO mutant WT plants.** (EXCEL)

## References

1. Lacroix C, Jolles A, Seabloom EW, Power AG, Mitchell CE, Borer ET. Non-random biodiversity loss underlies predictable increases in viral disease prevalence. Journal of the Royal Society Interface 2014; 11: 20130947. https://doi.org/10.1098/rsif.2013.0947.

2. Roossinck MJ. Lifestyles of plant viruses. Philosophical Transactions of the Royal Society B: Biological Sciences 2010; 365: 1899–1905. https://doi.org/10.1098/rstb.2010.0057.

3. Stobbe A, Roossinck MJ. Plant virus diversity and evolution. In: Wang A, Zhou X, editors. Current Research Topics in Plant Virology. Springer International Publishing; 2016. pp. 197–215. https://doi.org/10.1007/978-3-319-32919-2_8.

4. Elena SF, Agudelo-Romero P, Lalić J. The evolution of viruses in multi-host fitness landscapes. Open Virology Journal 2009; 3: 1–6. https://doi.org/10.2174/1874357900903010001.

5. Woolhouse MEJ. Population biology of multihost pathogens. Science 2001; 292: 1109–1112. https://doi.org/10.1126/science.1059026.

6. Whitlock MC. The Red Queen beats the jack-of-all-trades: the limitations on the evolution of phenotypic plasticity and niche breadth. American Naturalist 1996; 148: S65–S77. https://doi.org/10.1086/285902.

7. Lalić J, Cuevas JM, Elena SF. Effect of host species on the distribution of mutational fitness effects for an RNA virus. PLoS Genetics 2011; 11: e1002378.https://10.1371/journal.pgen.1002378.

8. Turner PE, Elena SF. Cost of host radiation in an RNA virus. Genetics 2000; 156, 1465–1470.

9. Bedhomme S, Lafforgue G, Elena SF. Multihost experimental evolution of a plant RNA virus reveals local adaptation and host-specific mutations. Molecular Biology and Evolution 2012; 29: 1481–1492. https://doi.org/10.1093/molbev/msr314.

10. Cervera H, Lalić J, Elena SF. Efficient escape from local optima in a highly rugged fitness landscape by evolving RNA virus populations. Proceedings of the Royal Society B: Biological Sciences 2016; 283: 1836. https://10.1098/rspb.2016.0984.

11. Bono LM, Draghi JA, Turner PE. Evolvability costs of niche expansion. Trends in Genetics 2020; 36: 14–23. https://doi.org/10.1016/j.tig.2019.10.003.

12. French RK, Holmes EC. An ecosystems perspective on virus evolution and emergence. Trends in Microbiology 2020; 28: 165–175. https://doi.org/10.1016/j.tim.2019.10.010.

13. Malmstrom CM, Hughes CC, Newton LA, Stoner CJ. Virus infection in remnant native bunchgrasses from invaded California grasslands. New Phytologist 2005; 168: 217–230. https://doi.org/10.1111/j.1469-8137.2005.01479.x.

14. Reusch TBH, Ehlers A, Hammerli A, Worm B. Ecosystem recovery after climatic extremes enhanced by genotypic diversity. Proceedings of the National Academy of Sciences of the United States of America 2005; 102: 2826–2831. https://doi.org/10.1073/pnas.0500008102.

15. Bedhomme S, Hillung J, Elena SF. Emerging viruses: Why they are not jacks of all trades? Current Opinion in Virology 2015; 10: 1–6. https://doi.org/10.1016/j.coviro.2014.10.006.

16. Woolhouse MEJ. Population biology of emerging and re-emerging pathogens. Trends in Microbiology 2002; 10: S3–S7. https://doi.org/10.1016/S0966-842X(02)02428-9.

17. Anderson PK, Cunningham AA, Patel NG, Morales FJ, Epstein PR, Daszak, P. Emerging infectious diseases of plants: Pathogen pollution, climate change and agrotechnology drivers. Trends in Ecology and Evolution 2004; 19: 535–544. https://doi.org/10.1016/j.tree.2004.07.021.

18. Cleaveland S, Haydon DT, Taylor L. Overviews of pathogen emergence: which pathogens emerge, when and why? In JE Childs, JS Mackenzie, JA Richt (Eds.), Wildlife and emerging zoonotic diseases: the biology, circumstances and consequences of cross-species transmission (pp. 85–111). Berlin, Germany: Springer 2007. https://doi.org/10.1007/978-3-540-70962-6_5.

19. Holmes EC. The evolutionary genetics of emerging viruses. Annual Review of Ecology, Evolution, and Systematics 2009; 40: 353–372. https://doi.org/10.1146/annurev.ecolsys.110308.120248.

20. Roossinck MJ, García-Arenal F. Ecosystem simplification, biodiversity loss and plant virus emergence. Current Opinion in Virology 2015; 10: 56–62. https://doi.org/10.1016/j.coviro.2015.01.005.

21. Bush WS, Moore JH. Genome-wide association studies. PLoS Computational Biology 2012; 8: e1002822. https://doi.org/10.1371/journal.pcbi.1002822.

22. Cantor RM, Lange K, Sinsheimer JS. Prioritizing GWAS results: a review of statistical methods and recommendations for their application. The American Journal of Human Genetics 2010; 86: 6–22. https://doi.org/10.1016/j.ajhg.2009.11.01.

23. Korte A, Farlow A. The advantages and limitations of trait analysis with GWAS: A review. Plant Methods 2013; 9: 29. https://doi.org/10.1186/1746-4811-9-29.

24. Zaitlen N, Kraft P. Heritability in the genome-wide association era. Human Genetics 2012; 131: 1655–1664. https://doi.org/10.1007/s00439-012-1199-6

25. Ohshima K, Yamaguchi Y, Hirota R, Hamamoto T, Tomimura K, Tan Z, et al. Molecular evolution of *Turnip mosaic virus:* evidence of host adaptation, genetic recombination and geographical spread. Journal of General Virology 2002; 83: 1511–1521. https://doi.org/10.1099/0022-1317-83-6-1511.

26. Yasaka R, Fukagawa H, Ikematsu M, Soda H, Korkmaz S, Golnaraghi A, et al. The timescale of emergence and spread of turnip mosaic potyvirus. Sci Rep 2017; 7: 4240. https://doi.org/10.1038/s41598-017-01934-7.

27. Navarro R, Ambrós S, Martínez F, Wu B, Carrasco JL, Elena SF. Defects in plant immunity modulate the rates and patterns of RNA virus evolution. bioRxiv 2020; 337402v1. https://www.biorxiv.org/content/10.1101/2020.10.13.337402v1.full.

28. Blüthgen N, Menzel F, Blüthgen N. Measuring specialization in species interaction networks. BMC Ecology 2006; 6: 9. https://doi.org/10.1186/1472-6785-6-9.

29. Hillung J, Cuevas JM, Valverde S, Elena SF. Experimental evolution of an emerging plant virus in host genotypes that differ in their susceptibility to infection. Evolution 2014; 68: 2467–2480. https://doi.org/10.1111/evo.12458.

30. González R, Butković A, Elena SF. Role of host genetic diversity for susceptibility-to-infection in the evolution of virulence of a plant virus. Virus Evolution 2019; 5: vez024. https://doi.org/10.1093/ve/vez024.

31. Pagán I, Fraile A, Fernandez-Fueyo E, Montes N, Alonso-Blanco C, García-Arenal F. *Arabidopsis thaliana* as a model for the study of plant–virus co-evolution. Philosophical Transactions of the Royal Society B: Biological Sciences 2010; 365: 1983–1995. https://doi.org/10.1098/rstb.2010.0062

32. 1001 Genomes Consortium. 1,135 genomes reveal the global pattern of polymorphism in *Arabidopsis thaliana*. Cell 2016; 166: 481–491. https://doi.org/10.1016/j.cell.2016.05.063.

33. Boyes DC. Growth stage-based phenotypic analysis of *Arabidopsis:* a model for high throughput functional genomics in plants. Plant Cell 2001; 13: 1449–510. https://doi.org/10.1105/tpc.010011.

34. Simko I, Piepho HP. The area under the disease progress stairs: calculation, advantage, and application. Phytopathology 2012; 102: 381–389. https://doi.org/10.1094/PHYTO-07-11-0216.

35. Corrêa RL, Sanz-Carbonell A, Kogej Z, Müller SY, Ambrós S, López-Gomollón S, et al. Viral fitness determines the magnitude of transcriptomic and epigenomic reprograming of defense responses in plants. Molecular Biology and Evolution 2020; 37: 1866–1881. https://doi.org/10.1093/molbev/msaa091.

36. Kone N, Asare-Bediako E, Silue S, Kone D, Koita O, Menzel W, et al. Influence of planting date on incidence and severity of viral disease on cucurbits under field condition. Annals of Agricultural Sciences 2017; 62: 99–104. https://doi.org/10.1016/j.aoas.2017.05.005.

37. Lippert C, Casale FP, Rakitsch B, Stegle O. LIMIX: Genetic analysis of multiple traits. bioRxiv 2014; 003905v2. https://www.biorxiv.org/content/10.1101/003905v2.

38. Seren Ü. GWA-Portal: Genome-wide association studies made easy. In D Ristova and E Barbez, editors. Root Development. New York: Springer; 2018. pp. 303–319. https://doi.org/10.1007/978-1-4939-7747-5_22.

39. Yin L, Zhang H, Tang Z, Xu J, Yin D, Zhang Z, et al. rMVP: A Memory-efficient, visualization-enhanced, and parallel-accelerated tool for genome-wide association study. bioRxiv 2020; 258491v1. https://www.biorxiv.org/content/10.1101/2020.08.20.258491v1.

40. Zhou X, Carbonetto P, Stephens M. Polygenic modeling with bayesian sparse linear mixed models. PLoS Genetics 2013; 9: e1003264. https://doi.org/10.1371/journal.pgen.1003264.

41. Zhou X, Stephens M. Genome-wide efficient mixed-model analysis for association studies. Nature Genetics 2012; 44: 821–824. https://doi.org/10.1038/ng.2310.

42. Makowski D, Ben-Shachar M, Lüdecke D. bayestestR: Describing effects and their uncertainty, existence and significance within the Bayesian framework. Journal of Open Source Software 2019; 4: 1541. https://doi.org/10.21105/joss.01541.

43. Ge X, Xia Y. The role of AtNUDT7, a Nudix hydrolase, in the plant defense response. Plant Signaling & Behavior 2008; 3: 119–120. https://doi.org/10.4161/psb.3.2.5019.

44. Dufresne PJ, Ubalijoro E, Fortin MG, Laliberté JF. *Arabidopsis thaliana* class II poly(A)-binding proteins are required for efficient multiplication of *Turnip mosaic virus*. Journal of General Virology 2008; 89: 2339–2348. https://doi.org/10.1099/vir.0.2008/002139-0.

45. Zhang T, Liu P, Zhong K, Zhang F, Xu M, He L, et al. *Wheat yellow mosaic virus* NIb interacting with host light induced protein (LIP) facilitates its infection through perturbing the abscisic acid pathway in wheat. Biology 2019; 8: 80. https://doi.org/10.3390/biology8040080.

46. Meyers BC, Kozik A, Griego A, Kuang H, Michelmore RW. Genome-wide analysis of NBS-LRR-encoding genes in *Arabidopsis*. Plant Cell 2003; 15: 809–834. https://doi.org/10.1105/tpc.009308.

47. Marone D, Russo M, Laidò G, De Leonardis A, Mastrangelo A. Plant nucleotide binding site–leucine-rich repeat (NBS-LRR) genes: active guardians in host defense responses. International Journal of Molecular Sciences 2013; 14: 7302–7326. https://doi.org/10.3390/ijms14047302.

48. Kant R, Tyagi K, Ghosh S, Jha G. Host alternative NADH: ubiquinone oxidoreductase serves as a susceptibility factor to promote pathogenesis of *Rhizoctonia solani* in plants. Phytopathology 2019; 109: 1741–1750. https://doi.org/10.1094/PHYTO-02-19-0055-R.

49. Ji DL, Lin H, Chi W, Zhang LX. CpLEPA is critical for chloroplast protein synthesis under suboptimal conditions in *Arabidopsis thaliana*. PLoS ONE 2012; 7: e49746. https://doi.org/10.1371/journal.pone.0049746.

50. Li D, Wei T, Abbott CM, Harrich D. The unexpected roles of eukaryotic translation elongation factors in RNA virus replication and pathogenesis. Microbiology and Molecular Biology Reviews 2013; 77: 253–266. https://doi.org/10.1128/MMBR.00059-12.

51. Sanfaçon H. Plant translation factors and virus resistance. Viruses 2015; 7: 3392–3419. https://doi.org/10.3390/v7072778.

52. Yoshimura K, Shigeoka S. Versatile physiological functions of the Nudix hydrolase family in *Arabidopsis*. Bioscience, Biotechnology, and Biochemistry 2015; 79:354–366. https://doi.org/10.1080/09168451.2014.987207.

53. Raja P, Sanville BC, Buchmann RC, Bisaro DM. Viral genome methylation as an epigenetic defense against geminiviruses. Journal of Virology 2008; 82: 8997–9007. https://doi.org/10.1128/JVI.00719-08.

54. McHale L, Tan X, Koehl P, Michelmore RW. Plant NBS-LRR proteins: Adaptable guards. Genome Biology 2006; 7: 212. https://doi.org/10.1186/gb-2006-7-4-212.

55. Lozano-Durán R, Rosas-Díaz T, Gusmaroli G, Luna AP, Taconnat L, Deng XW, et al. Geminiviruses subvert ubiquitination by altering CSN-mediated derubylation of SCF E3 ligase complexes and inhibit jasmonate signaling in *Arabidopsis thaliana*. Plant Cell 2011; 23: 1014–1032. https://doi.org/10.1105/tpc.110.080267.

56. Wu X, Ye J. Manipulation of jasmonate signaling by plant viruses and their insect vectors. Viruses 2020; 12: 148. https://doi.org/10.3390/v12020148.

57. Hillung J, García-García F, Dopazo J, Cuevas JM, Elena SF. The transcriptomics of an experimentally evolved plant-virus interaction. Scientific Reports 2016; 6: 24901. https://doi.org/10.1038/srep24901.

58. Rubio B, Cosson P, Caballero M, Revers F, Bergelson J, Roux F, Schurdi-Levraud V. Genome-wide association study reveals new loci involved in *Arabidopsis thaliana* and *Turnip mosaic virus* (TuMV) interactions in the field. New Phytologist 2019; 221: 2026–2038. https://doi/10.1111/nph.15507.

59. Hily JM, Poulicard N, Mora MÁ, Pagán I, García-Arenal F. Environment and host genotype determine the outcome of a plant-virus interaction: from antagonism to mutualism. New Phytologist 2016; 209: 812–822. https://doi.org/10.1111/nph.13631.

60. Xu P, Chen F, Mannas JP, Feldman T, Sumner LW, Roossinck MJ. Virus infection improves drought tolerance. New Phytologist 2008; 180: 911–921. https://doi.org/10.1111/j.1469-8137.2008.02627.x.

61. Chen CC, Chao CH, Chen CC, Yeh SD, Tsai HT, Chang CA. Identification of Turnip mosaic virus isolates causing yellow stripe and spot on calla lily. Plant Disease 2003; 87: 901–905. https://doi.org/10.1094/PDIS.2003.87.8.901.

